# Parasitism and host dispersal plasticity in an aquatic model system

**DOI:** 10.1101/2020.07.30.228742

**Authors:** Giacomo Zilio, Louise S. Nørgaard, Giovanni Petrucci, Nathalie Zeballos, Claire Gougat-Barbera, Emanuel A. Fronhofer, Oliver Kaltz

## Abstract

Dispersal is a central determinant of spatial dynamics in communities and ecosystems, and various ecological factors can shape the evolution of constitutive and plastic dispersal behaviours. One important driver of dispersal plasticity is the biotic environment. Parasites, for example, influence the internal condition of infected hosts and define external patch quality. Thus state-dependent dispersal may be determined by infection status and context-dependent dispersal by the abundance of infected hosts in the population. A prerequisite for such dispersal plasticity to evolve is a genetic basis on which natural selection can act. Using interconnected microcosms, we investigated dispersal in experimental populations of the freshwater protist *Paramecium caudatum* in response to the bacterial parasite *Holospora undulata*. For a collection of 20 natural host strains, we found substantial variation in constitutive dispersal, and to a lesser degree in dispersal plasticity. First, infection tended to increase or decrease dispersal relative to uninfected controls, depending on strain identity, potentially indicative of state-dependent dispersal plasticity. Infection additionally decreased host swimming speed compared to the uninfected counterparts. Second, for certain strains, there was a weak negative association between dispersal and infection prevalence, such that uninfected hosts tended to disperse less when infection was more frequent in the population, indicating context-dependent dispersal plasticity. Future experiments may test whether the observed differences in dispersal plasticity are sufficiently strong to react to natural selection. The evolution of dispersal plasticity as a strategy to mitigate parasite effects spatially may have important implications for epidemiological dynamics.

## Introduction

Dispersal, broadly defined as the movement of individuals with consequences for gene flow, is a key life-history trait (Bonte & Dahirel, 2017) driving metapopulation and metacommunity dynamics as well as the geographic distribution of species (Hanski, 1999). In recent years, the study of dispersal and dispersal syndromes have received increasing interest (Clobert *et al*., 2012; Stevens *et al*., 2014), as landscapes are seeing large-scale environmental alterations and fragmentation, rendering dispersal crucial to potentially mitigate these changes (Parmesan & Yohe, 2003; Cote *et al*., 2017). Although dispersal is often considered a constitutive trait, plastic dispersal behaviour represents a flexible alternative, responding to changes in the internal condition of an individual (state-dependent dispersal) and to external environmental factors (context-dependent dispersal) (Clobert *et al*., 2009). State-dependent dispersal has been associated with variation in factors such as body size, the developmental stage or sex of individuals (Bowler & Benton, 2005). In contrast, context-dependent dispersal decisions may be based on cues that provide information on biotic and abiotic patch properties, such as food availability, population density, or kin competition (see Ronce, 2007 and references therein).

In communities, dispersal plasticity may be advantageous in mitigating adverse interactions with other species (Fronhofer *et al*., 2015a). Parasites are particularly interesting in this respect: they are ubiquitous and impose strong selection pressures, and potentially drive the evolution of both state-dependent and context-dependent dispersal of their hosts (Iritani & Iwasa, 2014; Iritani, 2015; Narayanan *et al*., 2020; Deshpande *et al*., 2021). Empirical studies have investigated aspects of parasite-related dispersal (see below), but still little is known about the genetic basis of this kind of dispersal plasticity and its adaptive significance.

State-dependent dispersal may relate to morphological or physiological changes induced by parasites. The exploitation of host resources might decrease general activity levels, and thereby reduce movement and dispersal. Such negative effects have been documented for various organisms (Binning *et al*., 2017; Nørgaard *et al*., 2019; Baines *et al*., 2020), even though it is not necessarily a general rule (Nelson *et al*., 2015; Csata *et al*., 2017). While in many examples the observed effects may represent side effects, theory has identified conditions under which increased (but also decreased) dispersal when infected is adaptive, namely under kin selection (Iritani & Iwasa, 2014; Iritani, 2015) or when infection can be lost during dispersal (Shaw & Binning, 2016; Daversa *et al*., 2017). Indeed, increased dispersal of infected hosts is not uncommon (Suhonen *et al*., 2010; Brown *et al*., 2016), although it may also be the result of parasite manipulation (Lion *et al*., 2006; Martini *et al*., 2015).

Natural enemies may also produce context-dependent dispersal, as a means to reduce immediate predation or infection risk. For example, herbivores or predators can induce the production of specific dispersal morphs (Weisser *et al*., 1999; de la Pena *et al*., 2011). A recent multi-species study further showed that chemical predator-related cues increase dispersal probability (Fronhofer *et al*., 2018). Such cues may also exist in host-parasite systems, where infection-avoidance behaviour is well known (Behringer *et al*., 2006; Curtis, 2014). Recent theory shows that hosts may indeed evolve reaction norms, with dispersal being a function of the parasite infection prevalence (Deshpande *et al*., 2021). To date, few if any empirical studies have tested for the existence of such plastic population-level responses (French & Travis, 2001).

Adaptive phenotypic plasticity is a powerful solution in many situations (Chevin *et al*., 2013; Stamp & Hadfield, 2020), and just like constitutive traits, it has a genetic basis on which selection can act (Pigliucci, 2005; Garland & Kelly, 2006; Laitinen & Nikoloski, 2019). Dispersal-related traits have such a genetic basis (Saastamoinen *et al*., 2018) and constitutive dispersal can evolve rapidly in a parasite context (Koskella *et al*., 2011; Zilio *et al*., 2020). However, the genetics and evolution of dispersal plasticity is less well studied. In fact, how plastic dispersal varies between different genotypes under parasite challenge is rarely evaluated in empirical studies (Suhonen *et al*., 2010; Fellous *et al*., 2011), or the genetic diversity is treated as a random effect (Csata *et al*., 2017). Moreover, the number of genotypes evaluated is usually small, making it difficult to draw general conclusions (Leggett *et al*., 2013).

Here, using interconnected microcosms, we tested a collection of 20 natural strains of *Paramecium caudatum* for dispersal in the presence and absence of the bacterial parasite *Holospora undulata*. Previous work in this system had shown that infection reduces dispersal for a small number of strains (Fellous *et al*., 2011; Nørgaard *et al*., 2021). The first objective of the present study was to test whether this negative effect was general, or whether strains varied in infection-state dependent dispersal. Second, we tested for genetic variation in context-dependent dispersal by comparing the dispersal of uninfected hosts over a range of infection prevalences that had naturally established in the experimental populations. We found that parasite reduced or increased dispersal levels depending on strain identity, indicating a state-dependent plastic response of the infected hosts, but no general negative effect of infection. Furthermore, increasing infection prevalence tended to reduce host dispersal for certain strains, suggesting context-dependent dispersal plasticity of uninfected hosts. Such genetic variation in dispersal plasticity may provide the raw material for parasite-mediated selection, in natural settings or for the purpose of experimental evolution.

## Materials and methods

### Study system

*Paramecium caudatum* is a freshwater filter-feeding protist from stagnant waters of the Northern hemisphere (Wichterman, 2012). Like all ciliates, paramecia have a macronucleus for somatic gene expression and a germ-line micronucleus, used for sexual reproduction. The micronucleus can be infected by *Holospora undulata*, a gram-negative alpha-proteobacterium (Fokin, 2004). Infectious spores are released for horizontal transmission after host cell division or upon host death. Infectious spores are immobile and therefore rely on host movement or water current for their own dispersal. Vertical transmission occurs when hosts divide mitotically. Infection reduces *P. caudatum* division and survival (Restif & Kaltz, 2006) and also host dispersal (Fellous *et al*., 2011; Nørgaard *et al*., 2021).

### Experimental setup

#### Preparation of replicates

We established mass cultures for a collection of 20 genetically distinct strains of *P. caudatum* from different geographical regions (provided by S. Krenek, TU Dresden, Germany; Table S1, Supplementary Information). Distributed over two experimental blocks, 6 infected replicate cultures were established for each strain (20 strains x 2 blocks x 3 replicates = 120 replicates). Inocula were prepared from a mix of infected stock cultures in the lab, all originating from a single isolate of *H. undulata* established in 2001 (Dohra *et al*., 2013). Following standard protocols for the extraction of infectious spores (e.g., Nørgaard *et al*., 2021) we used c. 10^4^ spores to inoculate samples of c. 3-5 x 10^3^ host cells in 1.5 mL per assay replicate. Four days after inoculation, when infections have established, we expanded the cultures by regular addition of lettuce medium (supplemented with the food bacterium *Serratia marcescens*), until a volume of 50 mL was reached. In the same way, we set up three uninfected control populations per strain, giving a total of 180 experimental cultures. After three weeks, prior to the dispersal assay, population size (mean: 190 mL^-1^ ± 9 SE; 95% range [172; 208]) and infection prevalence (mean: 26.8 % ± 2.1; 95% range [3.1; 90.7]) had settled naturally in each experimental replicate.

#### Dispersal assay

We assayed the dispersal of infected and uninfected replicates in dispersal arenas, as described in Nørgaard *et al*. (2021). A dispersal arena consisted of three 50-mL Falcon tubes, linearly connected by 5-cm long silicon tubing (inner diameter: 0.8 cm). The 3-patch system was filled with 75 mL of medium to establish connections. Then the connections were blocked with clamps and 20 mL of a given replicate culture added into the middle tube. The lateral tubes received 20 mL of *Paramecium*-free medium. Connections were then opened, and the *Paramecium* allowed to disperse to the lateral tubes for 3h. After blocking the connections, we counted the individuals in samples from the middle tube (500-µl) and from the combined lateral tubes (3 mL) to estimate the number of non-dispersing and dispersing individuals (dissecting microscope, 40x). From the same samples, we also made lacto-aceto-orcein fixations (Görtz & Wiemann, 1989) and determined the infection status (infected / uninfected) of up to 30 dispersing and non-dispersing individuals, respectively (light microscope, phase contrast, 1000x). From the cell counts and the infection status data, we estimated the population density and infection prevalence in the middle tube at the beginning of the assay. From the same data, we also estimated the proportion of infected and uninfected dispersers for each replicate, referred to as per-3h “dispersal rate “ or dispersal, hereafter.

In addition, to investigate a potential link between dispersal and movement (Banerji *et al*., 2015; Pennekamp *et al*., 2019), we assayed swimming behaviour. For each strain, 1 infected and 1 uninfected individual were isolated from arbitrarily selected assay replicates, and allowed to replicate in a 2-mL plastic tubes for 8 days. For the resulting 40 monoclonal cultures (20 strains x 2 infection status) we placed 200-µL samples (10-20 individuals) on a microscope slide and recorded individual movement trajectories under a Perfex Pro 10 stereomicroscope, using a Perfex SC38800 camera (15 frames per second; duration: 10 s; total magnification: 10x). For each sample, average swimming speed (µm/s) and swimming tortuosity (standard deviation of the turning angle distribution, describing the extent of swimming trajectory change) were determined using video analysis (“BEMOVI “ package; Pennekamp *et al*., 2015).

### Statistical analysis

Statistical analyses were conducted in R, v. 3.6.3 (R Core Team, 2020) using Bayesian models with the ‘rstan ‘ (version 2.19.3) and ‘rethinking ‘ (version 2.0.1) packages (McElreath 2020).

For state-dependent dispersal, we compared the dispersal of the infected individuals (in infected replicates) with the dispersal in the uninfected control replicates. We fitted four models, from the intercept to the full interaction model, using a binomial regression with logit link function (chain length: warmup = 20,000 iterations, chain = 40,000 iterations). In the full model, the explanatory factors were infection status (infected or uninfected control), *Paramecium* strain identity, and the strain x status interaction. Experimental block only explained a negligible fraction of the dispersal variation (preliminary analysis, not shown) and was omitted from all further analyses. We fitted the models choosing vaguely informative priors; the intercepts and slope parameters followed a normal distribution with mean -2 and standard deviation 3 for the first, and mean 0 and standard deviation 1.75 for the latter. To account for overdispersion we included an observation-level random effect. The mean and standard deviation of the observation-level hyperprior followed a normal and half-normal distribution respectively, with mean 0 and standard deviation 1. The four state-dependent models were compared and ranked using the Watanabe-Akaike information criterion, WAIC (Watanabe, 2010), a generalized version of the Akaike information criterion (Gelman *et al*., 2014). The posterior predictions of the models were then averaged based on WAIC weights, and the relative importance (RI) of the explanatory variables was calculated as the sum of the respective WAIC model weights in which that variable was included. Due to loss of replicates, low population density, and/or very low levels of infection, 159 replicates (from 20 strains) of the 180 initial replicates were available for this analysis.

For context-dependent dispersal, we analysed the dispersal of uninfected *Paramecium* in infected assay replicates. We fitted 6 models, from the intercept to the full interaction model, using the same binomial regression with logit link function, chain lengths and prior specifications as above. The explanatory factors of the full model (varying intercept and slope) were infection prevalence, strain identity and the strain x infection prevalence interaction. The posterior predictions were averaged and ranked, and the RI calculated based on WAIC model weights as described above. For this analysis, 99 assay replicates (from 19 strains) of the initially 120 inoculated replicates were available.

We used similar analyses to test whether swimming speed and tortuosity varied as a function of strain identity and infection status. We standardized the response variable and fitted four models (from the intercept to the additive model, see Table S2 and S3, Supplementary Information) using a linear regression (chain length: warmup = 20,000 iterations, chain = 40,000 iterations) with an exponentially distributed prior (rate = 1) for standard deviation. As for the dispersal analysis, the parameter priors were vaguely informative; the intercept and slope parameters followed a normal distribution with mean 0 and standard deviation 2. We averaged and ranked the posterior predictions, and we obtained RI based on WAIC model weights. We further tested for correlations between these two swimming traits and mean strain dispersal, for infected and uninfected *Paramecium* (chain length: warmup = 2,000 iterations, chain = 10,000 iterations). Due to missing data, only 17 of the 20 strains were used for these analyses.

## Results

### State-dependent dispersal

Our analysis revealed substantial variation in constitutive dispersal among the 20 *P. caudatum* strains (relative importance, RI, of strain identity = 0.85; Table 1), ranging from 1% (95% compatibility Interval [0.001; 0.135]) to 41% ([0.02; 0.80]) of the individuals moving from the central to the lateral tubes (Fig. 1A). Our models provided limited evidence for state-dependent dispersal plasticity. Infection status (RI = 0.57) was retained in the best model fit (lowest WAIC; Model 3 in Table 1), indicating a general trend of infection to increase host dispersal. Even though the signal of the strain x infection status interaction (RI = 0.22) was only weak, patterns in Fig. 1B indicate that effects of infection varied with strain identity: several strains indeed dispersed more when infected (Fig. 1B right side of panel), but in at least half of the strains, infection had little effect or decreased host dispersal.

**Table 1.** Different statistical models and parameters included for the analysis and model averaging of the state-dependent dispersal. The rows represent the different models (the best model is highlighted in bold) and the columns the factors included in each model with the corresponding WAIC, standard error of the WAIC and WAIC weights. The RI row shows the relative importance of the explanatory variables.

|  | Strain | Status | Strain * Status | WAIC | SE | WAIC weight |
| --- | --- | --- | --- | --- | --- | --- |
| Model 1 |  |  |  | 1165 | 7.36 | 0.15 |
| Model 2 | X |  |  | 1163.8 | 27.76 | 0.27 |
| **Model 3** | **X** | **X** |  | **1163.3** | **27.67** | **0.35** |
| Model 4 | X | X | X | 1164.3 | 7.68 | 0.22 |
| RI | 0.85 | 0.57 | 0.22 |  |  |  |

**Figure 1.**
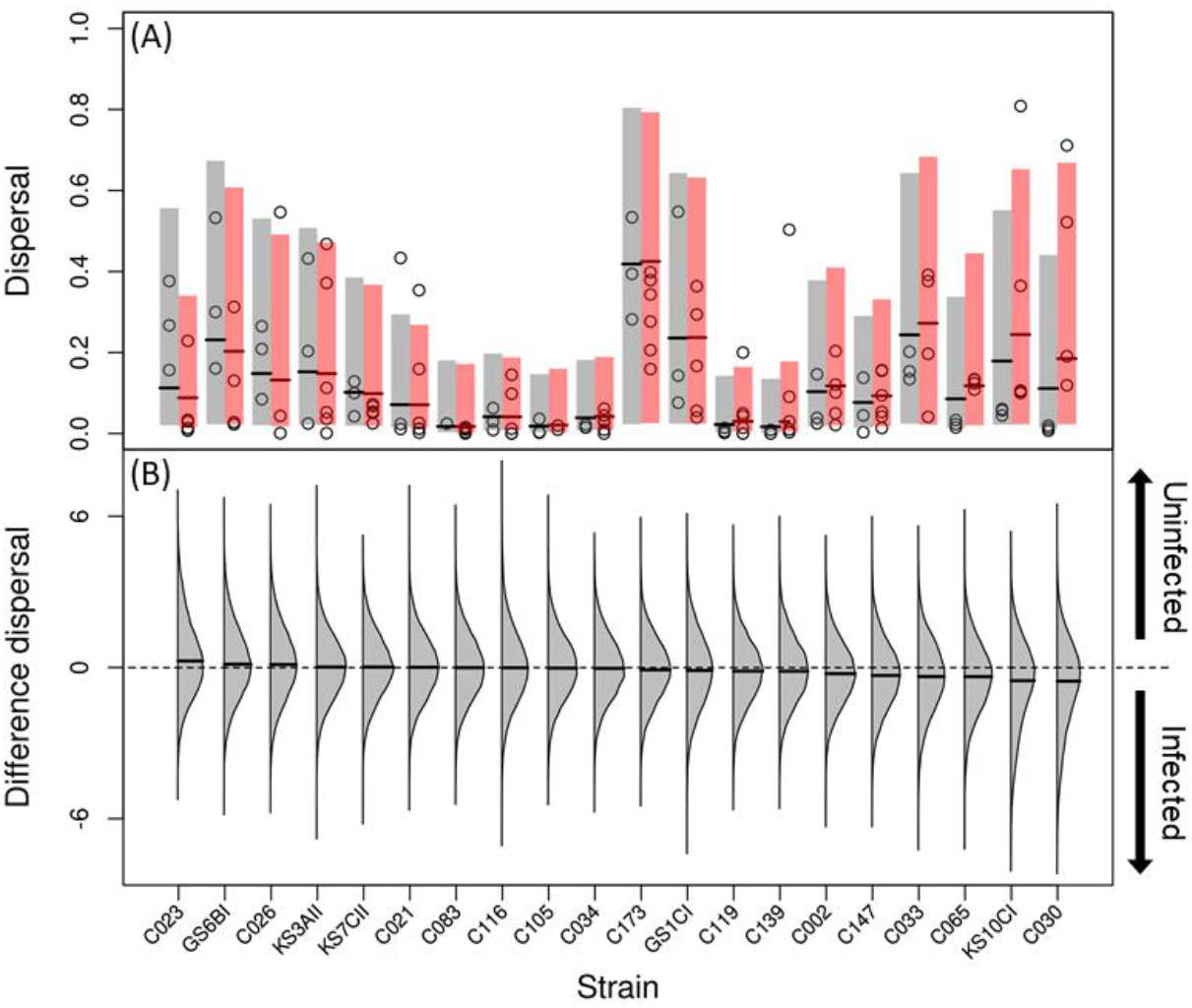
State-dependent dispersal of 20 *Paramecium caudatum* strains, as a function of infection status (uninfected / infected with *Holospora undulata)*. (A) Shaded bars and thick lines represent the 95% compatibility interval and the median of the averaged model predictions of the posterior distributions. Strains are ordered according to the difference between uninfected (grey) and infected (red) dispersal. Each circle represents an experimental replicate. (B) Difference between uninfected and infected averaged model posterior predictions for each strain (expressed in logits), the thick black line represents the median of the difference distribution. Distributions shifted below zero (dashed grey line) indicates higher dispersal in the infected (pointing-down arrow) compared to the uninfected (pointing-up arrow) treatment.

### Context-dependent dispersal

As in the above analysis, we found substantial genotypic variation in overall constitutive levels of dispersal for uninfected *Paramecium* (RI of strain identity = 0.80; Table 2). The best model (model 4 in Table 2) included an effect of infection prevalence (RI = 0.67), and thus context-dependent dispersal. Namely, uninfected individuals tended to disperse less at higher parasite infection prevalence in the population (Fig. 2): such negative dispersal-prevalence relationships were predicted for all but one strain (negative median slope values; Fig. 2B). To some degree, however, the strength of this relationship varied between strains (RI of infection prevalence x strain interaction = 0.21). As shown in Fig. 2B, distributions of predicted slopes show considerable variation and for the majority of strains there is considerable overlap with 0. Only a small number of strains (e.g., C139, C116, C083) show clearly negative slopes (Fig. 2B).

**Table 2.** Statistical models and parameters for the analysis and model averaging of the context-dependent dispersal. Each row represents a different model, the best model is highlighted in bold and the last row indicates the relative importance (RI) of the explanatory variables. The columns are the variables included in the six models with the corresponding WAIC, standard error of the WAIC and WAIC weights.

|  | Strain | Prevalence | Strain * Prevalence | WAIC | SE | WAIC weight |
| --- | --- | --- | --- | --- | --- | --- |
| Model 1 |  |  |  | 746.4 | 17.97 | 0.10 |
| Model 2 | X |  |  | 744.6 | 18.62 | 0.23 |
| Model 3 |  | X |  | 746.4 | 17.93 | 0.10 |
| **Model 4** | **X** | **X** |  | **744.2** | **18.77** | **0.28** |
| Model 5 | X | X |  | 746.7 | 18.16 | 0.08 |
| Model 6 | X | X | X | 744.8 | 18.61 | 0.21 |
| RI | 0.80 | 0.67 | 0.21 |  |  |  |

**Figure 2.**
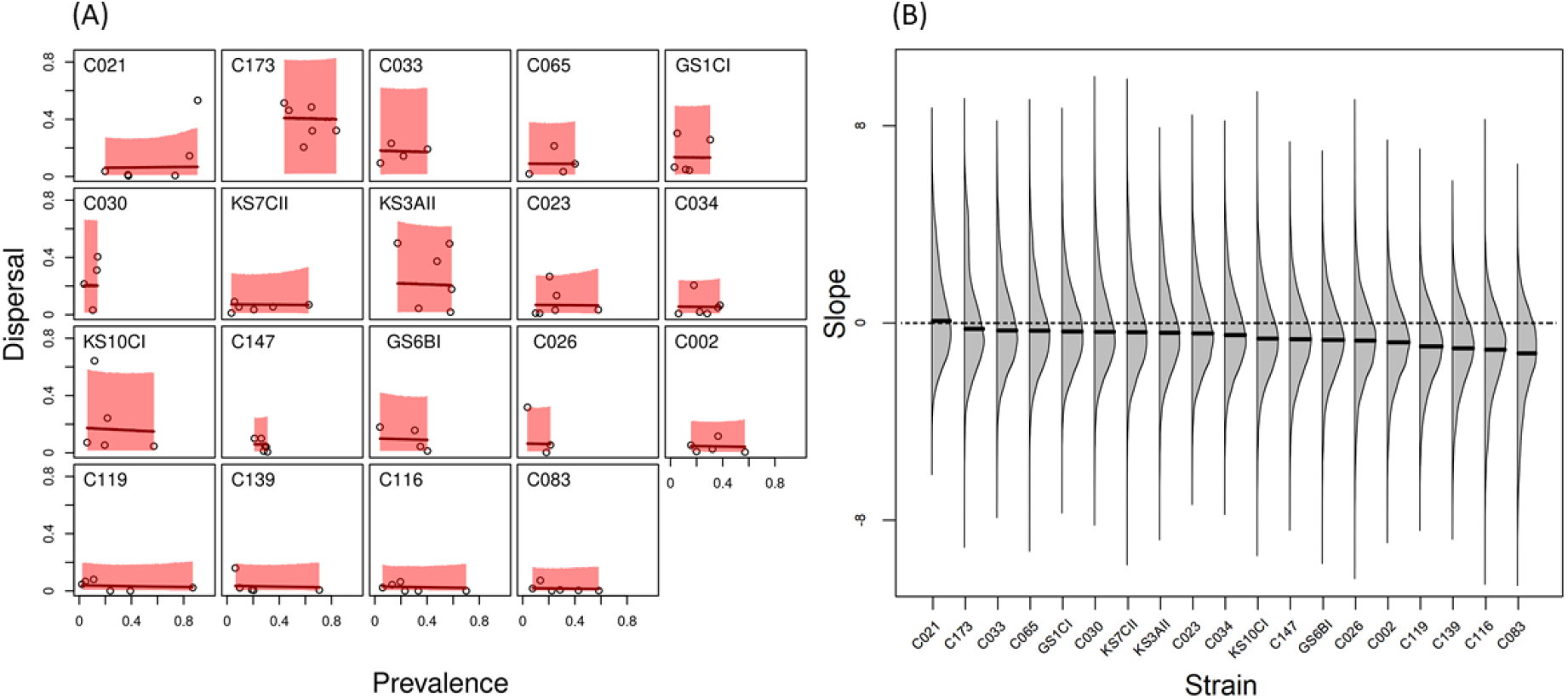
Context-dependent dispersal of 19 uninfected *Paramecium caudatum* strains, as a function of parasite (*Holospora undulata*) infection prevalence in the microcosm population. (A) Each panel represents a strain, and each circle an experimental replicate; the red shaded area and thick red lines are the 95% compatibility interval and median of the averaged model of the posterior distributions. (B) Averaging of the posterior distributions of the slope parameter calculated in logit (model 3-6, Table 2) with the thick black lines showing the median. Positive or negative slopes distributions (above or below zero, dashed grey line), indicate a higher or lower dispersal in response to increasing frequency of infected hosts.

### Swimming behaviour

The analysis of standardized swimming speed revealed strong effects of strain identity (RI = 0.9; Table S2) and infection status (RI = 1; Table S2). Namely, standardized swimming speed of uninfected *Paramecium* (median = 0.57, 95% CI [-0.64; 2.34]) was generally higher than that of infected ones (median = -1.20, 95% CI [-1.63; -0.77]), corresponding to a difference of almost 40% (median = 0.39, 95% CI [0.10; 0.68]; Fig. S1A-B). Swimming tortuosity was not affected by strain and weakly affected by infection status (RI strain = 0; RI status = 0.28; Table S3). Neither swimming speed (uninfected: r = 0.08, 95% CI [-0.39; 0.52]; infected: r = 0.07, 95% CI [-0.41; 0.54]) nor swimming tortuosity (uninfected: r = 0.15, 95% CI [-0.29; 0.56]; infected: r = -0.10, 95% CI [-0.55; 0.39]) were strongly correlated with dispersal.

## Discussion

Dispersal affects epidemiology and host-parasite (co)evolution in metapopulations (Lion & Gandon, 2015; Parratt *et al*., 2016), but how dispersal itself evolves due to antagonistic species interactions is less well known (Poethke *et al*., 2010; Drown *et al*., 2013; Deshpande *et al*., 2021). Here we focused on dispersal plasticity in response to parasitism, which may evolve as a means to reduce infection risk of the dispersing individuals and/or their relatives (Iritani & Iwasa, 2014; Iritani, 2015; Deshpande *et al*., 2021). Our study takes a first step towards an understanding of population-level processes, by measuring dispersal of infected and uninfected hosts in experimental microcosms and by exploring the genetic variation in plasticity for a collection of host strains. Overall, signals of dispersal plasticity were weak. Both infection status and infection prevalence modified dispersal to some degree, with at least some strains showing indications of state-dependent dispersal (i.e., when infected) and/or context-dependent dispersal (i.e., in response to infection prevalence).

### State-dependent plasticity: the dispersal of infected hosts

In previous studies, infection by *H. undulata* reduced dispersal in *P*. caudatum for a small set of strains (Fellous *et al*., 2011; Nørgaard *et al*., 2021). Here we used strains from a worldwide collection (Table S1) and find the entire range of trends, from negative or no impact of infection to even positive effects on host dispersal (Fig. 1). Reduced host dispersal may be explained by general negative effects of infection, through the energetic demand of an immune response, the diversion of host resources by the parasite or direct physical damage (Mideo, 2009). Indeed, *H. undulata* consumes nuclear proteins and nucleotides (Garushyants *et al*., 2018) and also causes massive interior swelling of the infected micronucleus, which would explain the clear and pervasive reduction in swimming speed observed in the complementary experiment (Fig. S1A-B). However, dispersal reductions were far from being universal, suggesting that the amount of host damage differs between genotypes. Differential fitness effects (virulence) and variation in resistance are known for this system (Restif & Kaltz, 2006), indicating the strong potential for genotypic-specificity in the responses to this parasite.

Moreover, it should be noted that the absence of a difference between infected and uninfected dispersal does not necessarily mean the absence of plasticity. Infected hosts may compensate parasite damage by re-allocating resources to maintain vital functions, such as foraging and feeding activity, and this may lead to a net-zero effect of infection on dispersal. Interestingly, some of our strains even seemed to ‘overcompensate ‘ and dispersed more when infected. Such a positive state-dependent dispersal may be selectively favoured in a metapopulation because it can reduce kin competition and kin infection (Iritani & Iwasa, 2014; Iritani, 2015; Deshpande *et al*., 2021). However, increased host dispersal may equally well reflect parasite manipulation, enhancing its dispersal to novel infection sites (Kamo & Boots, 2006; Lion *et al*., 2006; Martini *et al*., 2015).

The main purpose here was to quantify the (variation in) population-level effects of infection on dispersal. More work is needed to better understand the links between parasite action, host movement and dispersal. This concern, for example the relationship between parasite load, virulence and dispersal. Furthermore, unlike in other protists (Pennekamp *et al*., 2019), swimming speed was not a good predictor of dispersal. Other aspects of swimming behaviour (Ricci, 1989) may be more relevant in our system. Namely, *Paramecium* show a characteristic vertical distribution (Fels *et al*., 2008) relating to food and oxygen availability (Wichterman, 2012). Parasites are known to affect the position of hosts in the water column (Cezilly *et al*., 2000; Fels *et al*., 2004), and this may directly influence the probability of infected individuals finding the dispersal corridors in our microcosms.

### Context-dependent plasticity: the dispersal of uninfected hosts

Predator chemical signals induce dispersal in various organisms, including *P. caudatum* (Fronhofer et al. 2018). We tested for a similar parasite effect in our microcosm populations, by measuring the dispersal of uninfected hosts at different infection prevalences, with the assumption that higher prevalence equals a stronger signal of ‘parasite presence ‘. Unlike in the predator-cues study, we found little evidence for a positive dispersal-inducing effect. Dispersal decreased at higher infection prevalence, at least for certain strains. Interestingly, Deshpande *et al*. ‘s model (2021) predicts the evolution of such negative prevalence-dependent dispersal, as the result of complex spatio-temporal variations in eco-evolutionary processes. We do not know the evolutionary history of the strains, but our results suggest a possible genetic basis of context-dependent dispersal in this system and hence genetic variation that might be seen by natural selection.

Our experimental approach of using naturally established infection prevalence may not have produced strong enough signal variation for all strains. This could be remedied via more artificial designs, by mixing of infected and uninfected individuals to establish well-defined gradients. Infected cultures or inocula may also be filtered to specifically test for chemical cues (see Fronhofer *et al*., 2018). Finally, we made the simplifying assumption of linear dispersal reaction norms. However, dispersal responses may well follow non-linear rules, e.g., if there are signal thresholds (Fronhofer *et al*., 2015b), as observed for other traits (Morel-Journel *et al*., 2020) and predicted by Deshpande et al. (2021). Tests for non-linear relationships would require a much finer resolution (i.e., more replication) on the signal axis.

### Conditions for plasticity selection: outlook

The heritability of phenotypic plasticity of morphological or behavioural traits is generally lower than their constitutive heritability (Scheiner, 1993; Stirling *et al*., 2002). In line with this, we find much less among-strain differentiation for parasite-related dispersal plasticity than for constitutive dispersal, suggesting a weaker potential for responding to selection. However, the available genetic variation alone does not determine the relative importance of phenotypic plasticity in shaping evolutionary trajectories (Stamp & Hadfield, 2020). Phenotypic plasticity is generally favoured in variable, but nonetheless predictable environments (Leung *et al*., 2020). In a parasite context, dispersal plasticity evolution may thus depend on the spatio-temporal predictability of parasite encounter rates across a metapopulation (Deshpande *et al*., 2021). Additional factors are parasite virulence, the cost of dispersal (or its advantage if parasite release is possible during dispersal), or correlations with other traits (Iritani & Iwasa, 2014). For example, a recent experiment with the protist *Tetrahymena* revealed few genetic constraints on the concurrent evolution of plasticity across various traits (Morel-Journel *et al*., 2020). Indeed, state- and context-dependent dispersal might also evolve simultaneously in the presence of parasites, even though not necessarily in a correlated fashion (Deshpande et al., 2021). Our data indicate no genetic correlation between state- and context-dependent plasticity (r = -0.11, 95% CI [-0.55; 0.36]; based on strain averages), suggesting that independent responses to selection are possible, as shown in the model.

Our study represents one of the first accounts of the naturally existing genetic variation for state-dependent and context-dependent dispersal plasticity in relation to parasites. The signals of plasticity are weak and there are many open questions regarding the mechanistic and physiological basis of trait expression or information use. Nonetheless, in microbial systems such as ours, the observed variation opens promising avenues for future experiments. In microcosm landscapes, allowing the free interplay between dispersal and epidemiological processes, we can assess how dispersal plasticity affects parasite spread at the metapopulation level. Over longer time spans, we can also explore dispersal evolution and test evolutionary predictions on dispersal plasticity and its adaptive role in host-parasite interactions.

## Data availability statement

The experimental data will be made available upon potential acceptance (via Dryad/Figshare repository).

## Acknowledgements section

This work was supported by the Swiss National Science Foundation (grant no. P2NEP3_184489) to GZ, and by the 2019 Godfrey Hewitt mobility award granted to LN by ESEB, and by the. This is publication ISEM-XXXX-XXX of the Institut des Sciences de l‘Evolution.

## Supplementary Information

**Figure S1.**
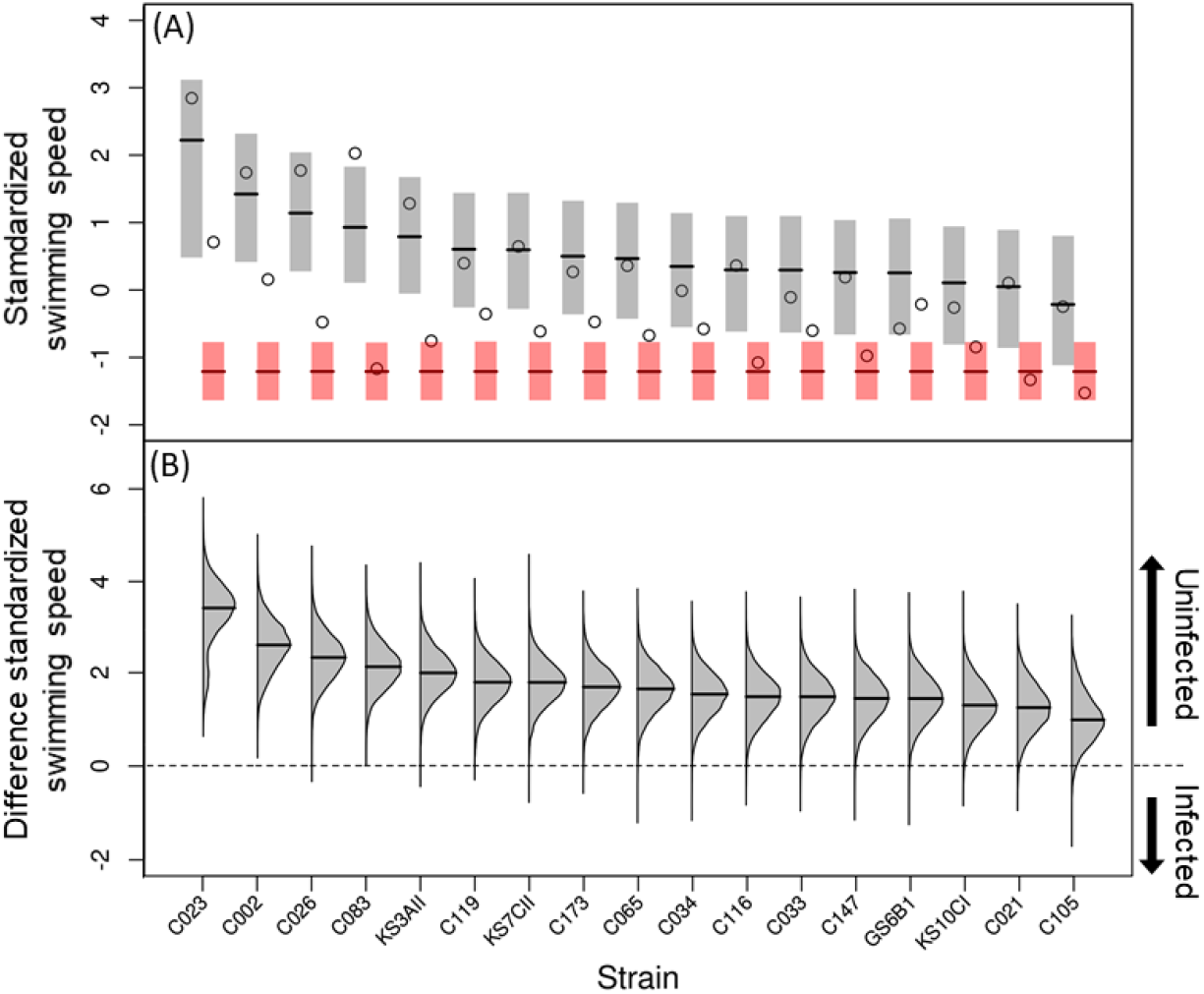
Standardized swimming speed of 17 *Paramecium caudatum*, as a function of infection status (uninfected / infected with *Holospora undulata)*. (A) Strains are ordered according to the difference between uninfected (grey) and infected (red) dispersal. Each circle represents an experimental replicate. Shaded bars and thick lines are the 95% compatibility interval and median of the averaged model predictions of the posterior distributions, and each circle represents the measured data of swimming speed per strain. (B) The difference in swimming speed between uninfected and infected averaged model posterior predictions for each strain. The thick black lines are the median of the difference distribution. Distributions shifted above zero (dashed grey line) indicates higher swimming speed in the uninfected treatment (pointing-up arrow) compared to the infected treatment (pointing-down arrow).

**Table S1.**
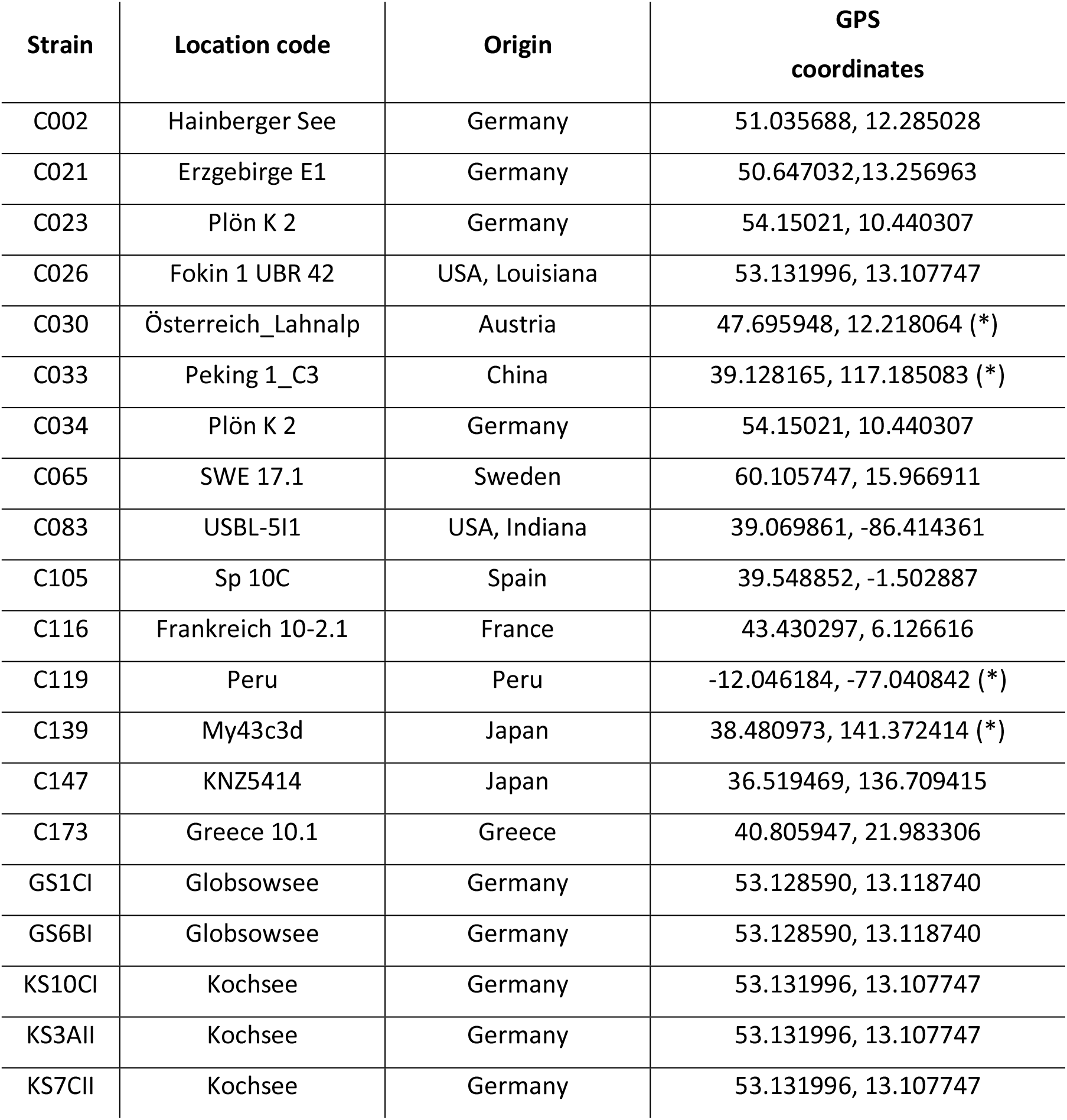
Host strain identity, location of origin and GPS coordinates, (*) indicates approximate location.

**Table S2.** Statistical models and parameters for the analysis and model averaging of the swimming speed. Each row shows a different model, from the intercept to the additive model. The best model is highlighted in bold and the last row indicates the relative importance (RI) of the strain and status effect. The columns are the explanatory variables included in the four models with the corresponding WAIC, standard error of the WAIC and WAIC weights.

|  | Strain | Status | WAIC | SE | WAIC weight |
| --- | --- | --- | --- | --- | --- |
| Model 1 |  |  | 100.0 | 10.45 | 0.0 |
| Model 2 | X |  | 114.1 | 5.84 | 0.0 |
| Model 3 |  | X | 83.7 | 10.29 | 0.1 |
| **Model 4** | **X** | **X** | **79.4** | **7.56** | **0.9** |
| **RI** | 0.9 | 1.0 |  |  |  |

**Table S3.**
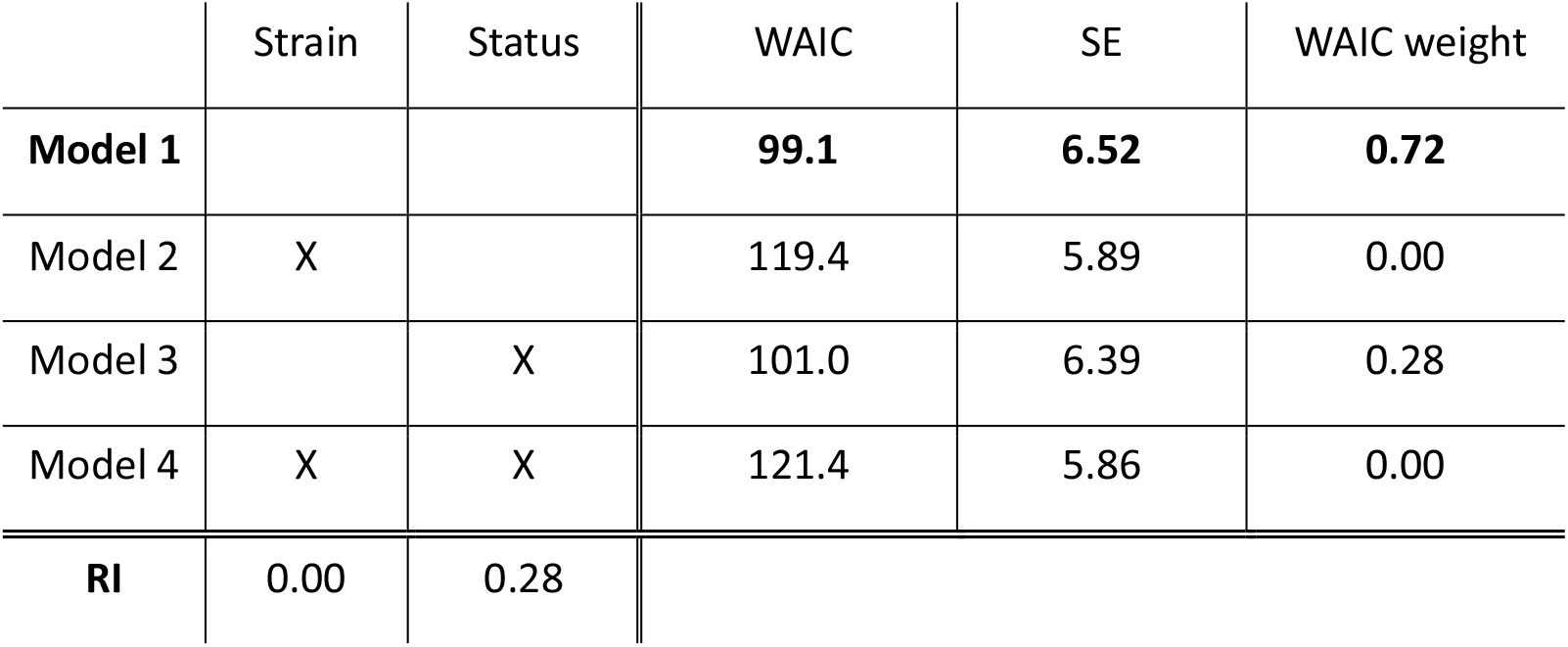
Models and parameters for the analysis and model averaging of the swimming tortuosity. Each row corresponds to a different model used for the analysis, with the best model is highlighted in bold. The last row shows the relative importance (RI) of the explanatory variables. The columns are the variables of the models with the corresponding WAIC, standard error of the WAIC and WAIC weights.

|  | Strain | Status | WAIC | SE | WAIC weight |
| --- | --- | --- | --- | --- | --- |
| **Model 1** |  |  | **99.1** | **6.52** | **0.72** |
| Model 2 | X |  | 119.4 | 5.89 | 0.00 |
| Model 3 |  | X | 101.0 | 6.39 | 0.28 |
| Model 4 | X | X | 121.4 | 5.86 | 0.00 |
| **RI** | 0.00 | 0.28 |  |  |  |

